# Variable paralog expression underlies phenotype variation

**DOI:** 10.1101/2022.04.27.489692

**Authors:** Raisa Bailon-Zambrano, Juliana Sucharov, Abigail Mumme-Monheit, Matthew Murry, Amanda Stenzel, Anthony T. Pulvino, Jennyfer M. Mitchell, Kathryn L. Colborn, James T. Nichols

## Abstract

Human faces are variable; we look different from one another. Craniofacial disorders further increase this variability. Here we used the zebrafish *mef2ca* mutant, which produces variable phenotypes, to understand craniofacial variation. Comparing different *mef2ca* alleles demonstrated that severity, measured by penetrance and expressivity, correlates with variation. Years of selective breeding for low and high penetrance produced strains that are either resilient, or sensitive, to the *mef2ca* mutation. Comparing these strains further demonstrates that severity correlates with variation. Gene expression studies indicated that selective breeding upregulated and downregulated *mef2ca* paralog expression in the low- and high-penetrance strains, respectively. We hypothesized that heritable paralog expression variation underlies mutant phenotype variation. In support, mutagenizing all *mef2ca* paralogs in the low-penetrance strain demonstrated modular buffering by paralogs. Specifically, some paralogs buffer severity while others buffer variability. We present a novel, mechanistic model for phenotypic variation where cryptic vestigial paralog expression modularly buffers development and contributes to evolution. These studies are a major step forward in understanding of the mechanisms of facial variation, including how some genetically resilient individuals can overcome a deleterious mutation.

## INTRODUCTION

### Human faces are variable

Human craniofacial variation allows us to identify each other. A recent study indicated that human facial structures are more variable than other human anatomical features, and that evolution selected for increased variation in craniofacial-associated genomic regions^1^. Thus, it is possible that high human facial variability is heritable, and under selection. While human genomic variation contributes to craniofacial phenotypic variation^2, 3^, facial variation persists among genetically homogeneous populations^1^. Therefore, even among individuals with similar genomes, craniofacial development may remain noisy, or sensitive to minor fluctuations, compared with other developmental processes. Penetrance is the frequency of a phenotype associated with a genotype.

Expressivity is different degrees of the same phenotype associated with a genotype. Here, we use both penetrance and average expressivity to measure phenotype severity. We also use the spread in expressivity to measure phenotypic variation. Further, there are two types of variation. Among-individual variation can be quantified by measuring differences between individuals. Within-individual variation is measured by quantifying departures from symmetry on left versus right sides of an individual, known as fluctuating asymmetry^4^. It is unknown if among- and within-individual variation are products of the same biological mechanisms^5^. There is evidence to support that these two types of variation are associated^6, 7^, however other studies indicate that they are independent^8, 9^. Further work is needed to resolve this question.

### Variability in human craniofacial disease

In the 1940’s Waddington observed that genetic mutations in model organisms are often associated with increased phenotypic variation compared to wild types^10^, and more recent studies support this finding^11–14^.

Human genetic craniofacial disease phenotypes also appear more variable than normal human facial phenotypes. Facial clefting among monozygotic twins can be incompletely penetrant^15, 16^, demonstrating among-individual variation. How the same genetic disease allele can have devastating consequences in some individuals while other resilient individuals can overcome the deleterious mutation is not well understood^17^.

One human genetic disease that presents variable craniofacial phenotypes is *MEF2C* haploinsufficiency syndrome. Patients with mutations affecting the transcription factor encoding gene *MEF2C* show variable facial dysmorphologies^18–20^. Additionally, craniofacial asymmetry is documented in some patients with this disorder^21, 22^. Thus, both among- and within-individual variation are present in *MEF2C* haploinsufficiency syndrome patients. Numerous *MEF2C* mutant alleles cause this disorder. Strikingly, one of the mildest documented cases of this disease is a large deletion encompassing the *MEF2C* locus^23^. It remains unknown if the among-individual variation in this disease is due to *MEF2C* allelic variation, and if full deletions are more mild than other alleles.

### Variable buffering by gene family members might underlie among-individual variation

During evolution, whole genome duplications produced multiple *mef2* genes, or paralogs, in vertebrate genomes. Thus far, none are associated with craniofacial development besides *mef2c*. When these duplications occur, the most likely outcome is loss of one of the duplicates through the accumulation of deleterious mutations and eventual nonfunctionalization^24, 25^. However, sometimes mutations occur in gene regulatory elements partitioning gene expression domains among the new duplicates. This preserves both copies and is called subfunctionalization^26^. Subfunctionalized duplicates may experience only partial loss of expression subdomains^26^, retaining relics of their ancestral expression pattern even if the gene is no longer required for the original function. We hypothesize that this vestigial expression buffers against loss of another paralog. To our knowledge, this vestigial buffering hypothesis has not been previously proposed or tested.

We previously found evidence to support this hypothesis in zebrafish. *mef2ca* single mutants produce dramatically variable craniofacial phenotypes^27^. *mef2cb* is the most closely related *mef2ca* paralog, but is not required for craniofacial development and homozygous mutants are viable^28^. However, we reported that ventral cartilage defects associated with *mef2ca* mutation become more severe when we removed a single functional copy of *mef2cb* from *mef2ca* homozygous mutants^29^. Our vestigial buffering hypothesis predicts that although *mef2cb* is no longer overtly required for craniofacial development, remaining *mef2cb* expression in neural crest cells might partially substitute for *mef2ca* loss. There is further evidence from mouse mutants that *Mef2* paralogs can functionally substitute for one another^30^. Because no other zebrafish *mef2* gene mutants have been reported, their function is unknown.

### We developed a system for understanding craniofacial development and variability

One prominent phenotype in zebrafish *mef2ca* homozygous mutants is expansion of the opercle, a bone supporting the gill flap. This phenotype is remarkably variable^29, 31^; *mef2ca* homozygous mutants have many opercle shapes and some even develop wild-type looking opercle bones^27^. With some genes, mutants encoding a premature termination codon (PTC) upregulate compensating genes through transcriptional adaptation, explaining why some mutant alleles do not produce a phenotype^32, 33^. Transcriptional adaptation does not contribute to incomplete penetrance in our system^34^. We demonstrated that the mechanisms underlying opercle variation are heritable through selective breeding, which shifted the penetrance of the expanded bone phenotype to generate strains with consistently low or high penetrance of this phenotype^27^. *mef2ca* is pleiotropic, and although we only selected on opercle bone penetrance, nearly all *mef2ca* associated phenotypes changed penetrance as well^34^.

The only phenotype remaining fully penetrant in the low- and high-penetrance strains is a shortened symplectic cartilage, a linear, rod-shaped cartilage that functions as a jaw support structure. While in unselected lines, *mef2ca*-associated phenotypes are only found in *mef2ca* homozygous mutants^35^, in high-penetrance heterozygotes we observed the shortened symplectic cartilage, suggesting this phenotype is the most sensitive to *mef2ca* loss^34^. However, penetrance is a binary measurement (a shortened symplectic is present or not), and we have not yet examined expressivity (to what extent a mutant symplectic is shortened). We do not know if the average length of the shortened symplectic, or the spread in symplectic length is different between low- and high-penetrance strains, or with different *mef2ca* mutant alleles.

Here we capitalize on the strengths of our zebrafish system to address such unanswered questions about craniofacial variation. We quantified expressivity by taking linear measurements of the symplectic cartilage, allowing us to measure severity, among-individual variation, and within-individual variation in the zebrafish craniofacial skeleton. Combining these measurements with penetrance scoring, we compared phenotype severity and variation in several *mef2ca* alleles, including a large deletion. We were able to order this allelic series according to severity, finding that more severe alleles have more among-individual variation, but not more within-individual variation. Comparing selectively bred strains, we found a more severe symplectic phenotype, and more among-individual variation, but not within-individual variation in the high-penetrance strain. We compared the expression of the *mef2* paralogs between selectively-bred strains and found that many of the paralogs are upregulated in the low-penetrance strain compared with the high-penetrance strain. To determine if the increased paralog expression we discovered in the low-penetrance strain buffers the *mef2ca* mutant phenotype, we mutagenized the *mef2* paralogs in the low-penetrance strain. Double mutant analyses indicate that the different paralogs modularly buffer different aspects of the pleiotropic *mef2ca* mutant phenotype, some affecting penetrance, while others affect expressivity. These findings demonstrate that heritable, variable paralog expression is a major factor affecting phenotype severity, and among-individual phenotypic variation, but does not contribute to within-individual variation.

## RESULTS

### Among-individual, but not within-individual, variation correlates with allele severity

The phenotypic variation often associated with human genetic diseases might be partially due to individuals inheriting different mutant alleles retaining different levels of functional activity. A strength of the zebrafish system is that the same mutant allele can be analyzed in many different individuals, and genetic background can be controlled. To determine how different mutant alleles might contribute to phenotypic variation in our system, we examined four different *mef2ca* alleles. The wild-type *mef2ca* allele encodes an N-terminal MADS box (MCM1, agamous, deficiens, SRF), and an adjacent MEF2 domain^36^ (Fig. 1A). These highly conserved domains mediate dimerization, DNA binding, and co-factor interactions^37^. The C-terminal domain is more divergent among different *mef2* genes^38^. The *mef2ca^b631^* mutant allele arose from a forward genetic screen and destroys the initiating methionine^35^. The *mef2ca^b1086^* mutant allele also arose from a forward genetic screen and produces a PTC just downstream of the MADS box. The final mutant allele, *mef2ca^co3008^*, was created for this study to model the full *MEF2C* deletion allele found in a mild patient^23^ (see Introduction). All the protein-coding exons and portions of the 5’ and 3’ untranslated regions are removed in this deletion allele. This range of alleles allows us to examine how different mutations produce phenotypes with different severity and variability.

**Figure 1:**
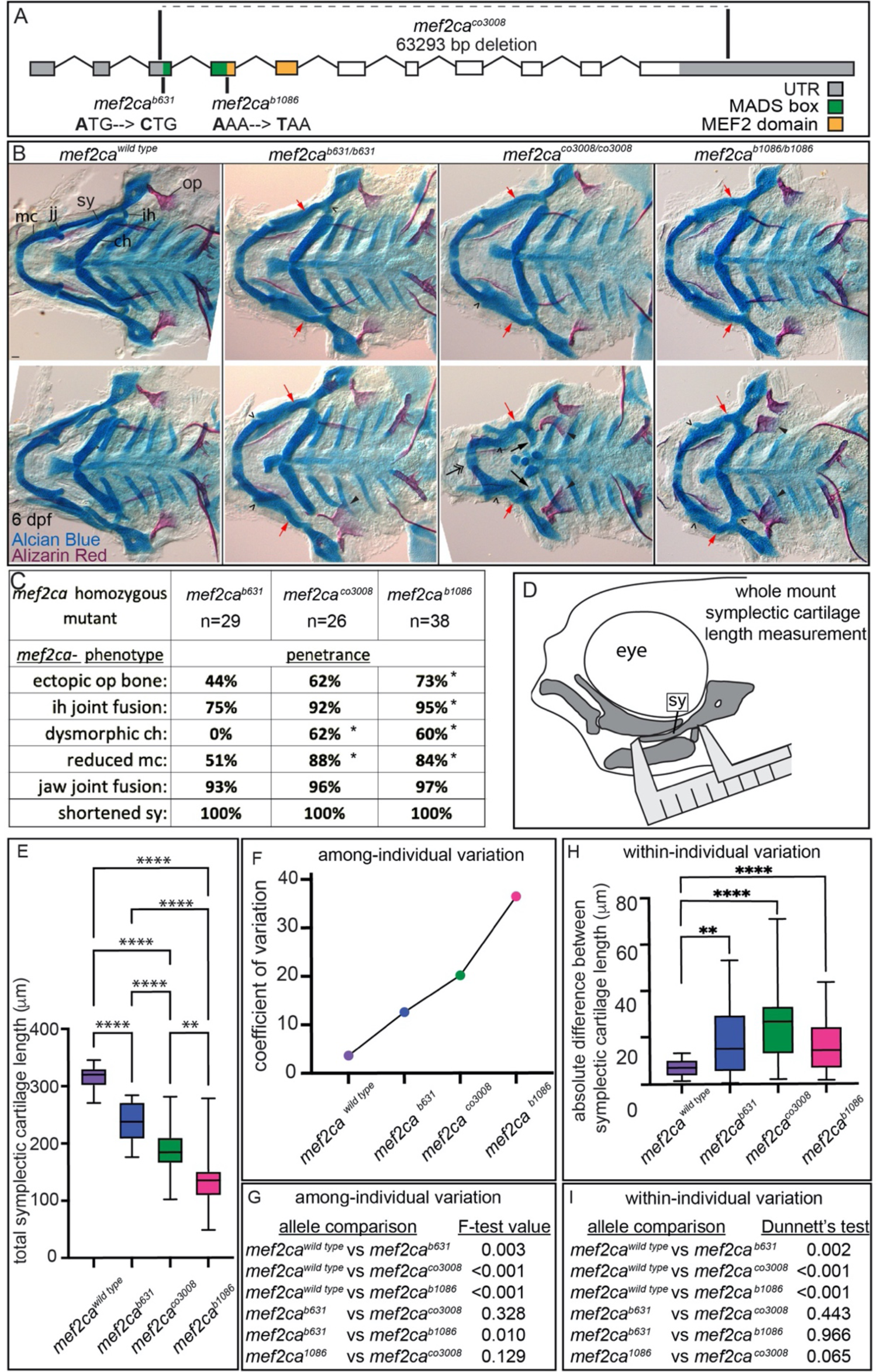
A *mef2ca* allelic series demonstrates that severity and among-individual variability are correlated. (A) Schematic of *mef2ca* exonic structure. Mutant alleles used in this study and regions encoding known functional domains are annotated. (B) Wild-type, or zebrafish heterozygous for the indicated *mef2ca* alleles, were pairwise intercrossed and six days post fertilization (dpf) larvae were stained with Alcian Blue and Alizarin Red to label cartilage and bone. The individuals were then genotyped, flat mounted, and imaged. Two examples (upper, lower) are provided for each genotype. The following craniofacial skeletal elements are indicated in a wild-type individual: opercle bone (op), branchiostegal ray (br), Meckel’s (mc), ceratohyal (ch), symplectic (sy) cartilages, interhyal (ih) and jaw (jj) joints. Indicated phenotypes associated with *mef2ca* mutants include: ectopic bone (arrowheads), interhyal and jaw-joint fusions (^), dysmorphic ch (arrows), reduced mc (double arrowhead), and a shortened sy (red arrows). Scale bar: 50μm (C) The penetrance of *mef2ca* mutant-associated phenotypes observed in 6 dpf larvae homozygous for each allele are indicated. Asterisk indicates significant difference in penetrance between the indicated allele and *mef2ca^b631^* by Fishers exact test. There are not significant penetrance differences between *mef2ca^co3008^* and *mef2ca^b1086^*. (D) Schematic indicating how the symplectic cartilage length was measured in this study. (E) Symplectic cartilage length was measured from 6 dpf larvae homozygous for the indicated *mef2ca* allele. The p-values from a Dunnet’s T3 test are indicated (** ≤ 0.01, **** ≤ 0.0001). (F) The coefficient of variation for symplectic length in fish homozygous for each allele was plotted. (G) Table listing F-test values testing for significant differences in variation between fish homozygous for each allele. (H) Symplectic cartilage length on left and right sides of 6 dpf zebrafish was measured to determine fluctuating asymmetry or the absolute difference between left and right for fish homozygous for the indicated alleles. There is no significant difference in fluctuating asymmetry between any of the mutant alleles. (I) Table listing the Dunnett’s test for significant differences in fluctuating asymmetry between fish homozygous for each allele. For box and whisker plots, the box extends from the 25th to 75th percentiles. The line in the middle of the box is plotted at the median and the bars are minimum and maximum values. N’s for all analyses are indicated in C.

All of the mutant alleles variably display the same *mef2ca* mutant-associated phenotypes including ectopic bone near the opercle, interhyal joint fusion, dysmorphic ceratohyal, reduced Meckel’s cartilage, jaw joint fusion, and shortened symplectic (Fig. 1B, C). Skeletal preparations from two full-sibling individuals illustrate both among-individual and within-individual variation easily observed in *mef2ca* mutants (Fig. 1B). Comparing the two *mef2ca^co3008^* individuals (upper and lower panels) demonstrates among-individual variation associated with the opercle bone phenotype; one individual has phenotypically wild-type opercles while the other individual has bilateral mutant phenotypes. Meanwhile, within-individual (left-right) opercle phenotype variation is present in one of the *mef2ca^b631^* animals (lower).

To understand the variation associated with these different alleles, we first scored penetrance of the various *mef2ca* phenotypes in each mutant allele. We determined that the frequency of most phenotypes was lowest in *mef2ca^b631^*, followed by *mef2ca^co3008^*, then *mef2ca^b1086^* (Fig. 1C). These results allow us to order this allelic series by penetrance; *mef2ca^b631^* is the mildest mutant allele and *mef2ca^b1086^* is the most severe. We next used the linear symplectic length to examine expressivity (Fig. 1D). The average symplectic length is significantly different between every pair of alleles. The *mef2ca* wild-type symplectic cartilage is the longest followed by *mef2ca^b631^*, *mef2ca^co3008^*, then *mef2ca^b1086^* is the shortest (Fig. 1E). Ordering our allelic series by expressivity shows the same order with respect to severity as the penetrance study (Fig. 1C). These findings also support our previous discovery that transcriptional adaptation^32,33^ is not a major factor underlying phenotypic variation in our system^34^; the PTC (*mef2ca^b1086^*) allele would be expected to be more mild than the full deletion (*mef2ca^co3008^*) allele if transcriptional adaptation was a factor.

Comparing the total (left side plus right side) symplectic cartilage length among full-sibling individuals allowed us to examine the among-individual variation associated with each allele. Strikingly, we found that variation is positively correlated with severity; the most severe allele is also the most variable (Fig. 1F). The phenotypes associated with all the mutant alleles were significantly more variable than the wild type, and the most severe mutant (*mef2ca^b1086^*) allele was significantly more variable than the mildest mutant (*mef2ca^b631^*) allele (Fig. 1G). When we examined within-individual variation by determining the absolute value of the difference between left and right symplectic cartilage length, we found that while all mutant alleles have more within-individual variation than the wild type, there is no significant difference between mutant alleles for this type of variation (Fig. 1H, I). Our finding that among-individual variation, but not within-individual variation correlates with severity suggests that, in our system, these two types of variation are mechanistically different and are buffered by different factors.

### Selective breeding reveals that among-individual, but not within-individual, variation is heritable and segregates with penetrance

Severity is positively correlated with variation when comparing different *mef2ca* alleles on the same AB genetic background. Therefore, we hypothesized that the same mutant allele, *mef2ca^b1086^*, maintained on two different genetic backgrounds might also reveal that among-individual variation is positively correlated with severity. An ongoing, long-term selective breeding experiment in our laboratory derived two strains of zebrafish with consistently low and high penetrance of *mef2ca-*associated phenotypes in *mef2ca^b1086^* homozygous mutants^27, 34, 39^ (Fig. 2A). Our previous work examined the phenotypes present in homozygous mutant fish from these strains^34^. Here, we closely examined *mef2ca* wild types (*mef2ca*^+/+^) from these strains. We were surprised to observe some *mef2ca*-associated phenotypes, like shortened symplectic cartilages and fused interhyal joints, were occasionally observed in *mef2ca*^+/+^ individuals from the high-penetrance strain (Fig. 2B). Importantly, these phenotypes are not likely due to general developmental delay because other skeletal structures like pharyngeal teeth are unaffected. Thus, the phenotypes we discovered in high-penetrance *mef2ca*^+/+^ individuals are specific to developmental processes associated with *mef2ca* function. Quantifying the severity of the shortened symplectic cartilage phenotype, we found significant differences between low- and high-penetrance strains for both *mef2ca*^+/+^, as well as *mef2ca*^+/-^ (Fig. 2C).

**Figure 2:**
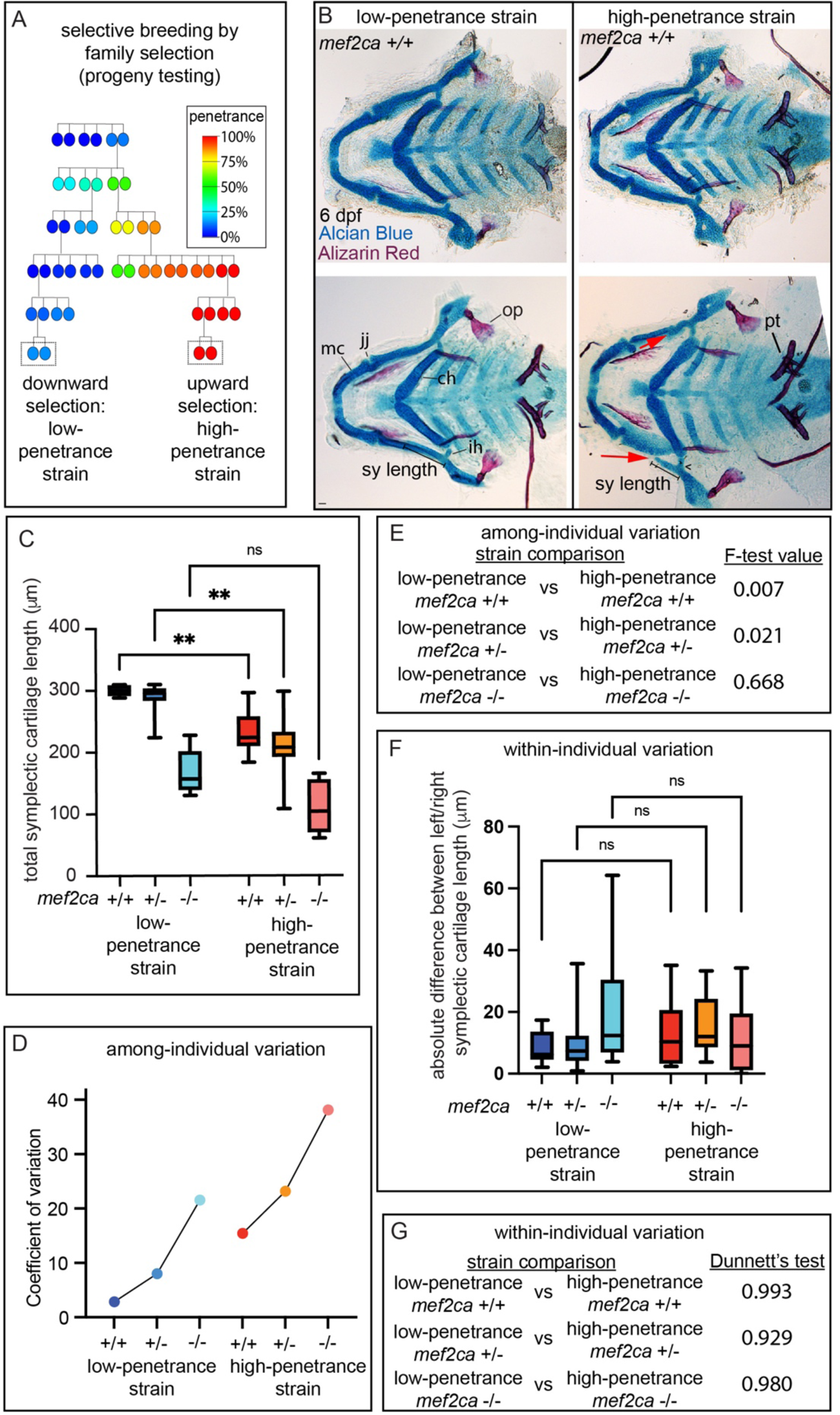
Selective breeding affects *mef2ca*-associated phenotype severity and variation in *mef2ca* wild types, heterozygotes, and homozygous mutants. (A) Selective breeding pedigree illustrating ectopic bone phenotype penetrance inheritance. Dashed boxes indicate families used in this study. (B) Alcian Blue and Alizarin Red stained animals from the low- and high-penetrance strains were genotyped, and *mef2ca* homozygous wild types were flat mounted and imaged. The following craniofacial skeletal elements are indicated in a wild-type individual from the low-penetrance strain: opercle bone (op), Meckel’s (mc), ceratohyal (ch), symplectic (sy) cartilages, interhyal (ih) and jaw (jj) joints. Phenotypes normally associated with *mef2ca* homozygous mutants are present in some wild types from the low-penetrance strain including: ih joint fusions (^), and shortened sy (red arrows). Bars indicating sy length are presented to illustrate the shortenend symplectic phenotype present in some high-penetrance wild types but not low-penetrance wild types. A stage-appropriate complement of ankylosed of pharyngeal teeth (pt) are present and normal sized op bones are present in the individual with shortened sy, indicating the phenotypes we discovered in high-penetrance *mef2ca^+/+^* are not due to general delay. Scale bar: 50μm (C) Symplectic cartilage length was measured from 6 dpf larvae from wild types, heterozygotes, and homozygous mutants from both the low- and high-penetrance strains. P values from a Dunnet’s T3 test are indicated (** ≤ 0.01) (D) The coefficient of variation for symplectic length in all three genotypes from both strains was plotted. (E) Table listing F-test values testing for significant differences in variation between strains comparing the same genotype. (F) Symplectic cartilage length on left and right sides of 6 dpf zebrafish was measured to determine fluctuating asymmetry, or the absolute difference between left and right for all three genotypes from both strains. (G) Table listing the Dunnett’s test for significant differences in fluctuating asymmetry between all three genotypes from both strains. For box and whisker plots, the box extends from the 25th to 75th percentiles. The line in the middle of the box is plotted at the median and the bars are minimum and maximum values.

When we examined variation, we observed that among-individual variation again positively correlated with severity (Fig. 2D). Even *mef2ca* homozygous wild types exhibited significantly higher severity and variation in the high-penetrance strain compared with the low-penetrance strain (Fig. 2C, E). Because the low- and high-penetrance strains are both similarly inbred, it is unlikely that there is more genetic variation in the high-penetrance strain background accounting for the increased variation in this strain.

Consistent with comparing different alleles (Fig.1), we found that among-individual variation correlated with severity while within-individual variation did not (Fig. 2F, G). These data provide further evidence that the two types of variation we examine here are regulated by different underlying mechanisms in our system. Moreover, it is fascinating that a phenotype originally associated with a homozygous loss of function mutation can appear in homozygous wild types after selective breeding for high penetrance. We previously observed a similar phenomenon with heterozygotes in the high-penetrance strain, and concluded that this is a form of genetic assimilation^34, 40^.

### Six *mef2* paralogs in the zebrafish genome share highly conserved amino acid sequences

Wild-type siblings of *mef2ca* mutants from the selectively bred high-penetrance strain display *mef2ca* mutant-associated phenotypes. One mechanistic hypothesis accounting for this discovery is that selective breeding changed the expression of another gene, or genes, which function similarly to *mef2ca* to generate a ‘natural knock down’ in *mef2ca* wild types. The five *mef2ca* paralogs are strong candidates for participating in *mef2ca*-directed developmental processes. We used Clustal Omega to compare the amino acid sequences of these paralogs, finding that the closest related paralog to *mef2ca* is *mef2cb.* Together these are the co-orthologs of mammalian *Mef2c* (Fig. 3A). *mef2d* is the next most closely related paralog to *mef2ca. mef2aa* and *mef2ab*, co-orthologs of mammalian *mef2a*, are next. The most distantly related family member is *mef2b*. There is only one zebrafish ortholog for *Mef2b* and *Mef2d,* presumably the other co-ortholog for each was lost. The zebrafish *mef2* paralogs each encode a MADS box and MEF2 domain which are remarkably similar (Fig. 3B and S1), suggesting that they may perform a similar function to *mef2ca*.

**Figure 3.**
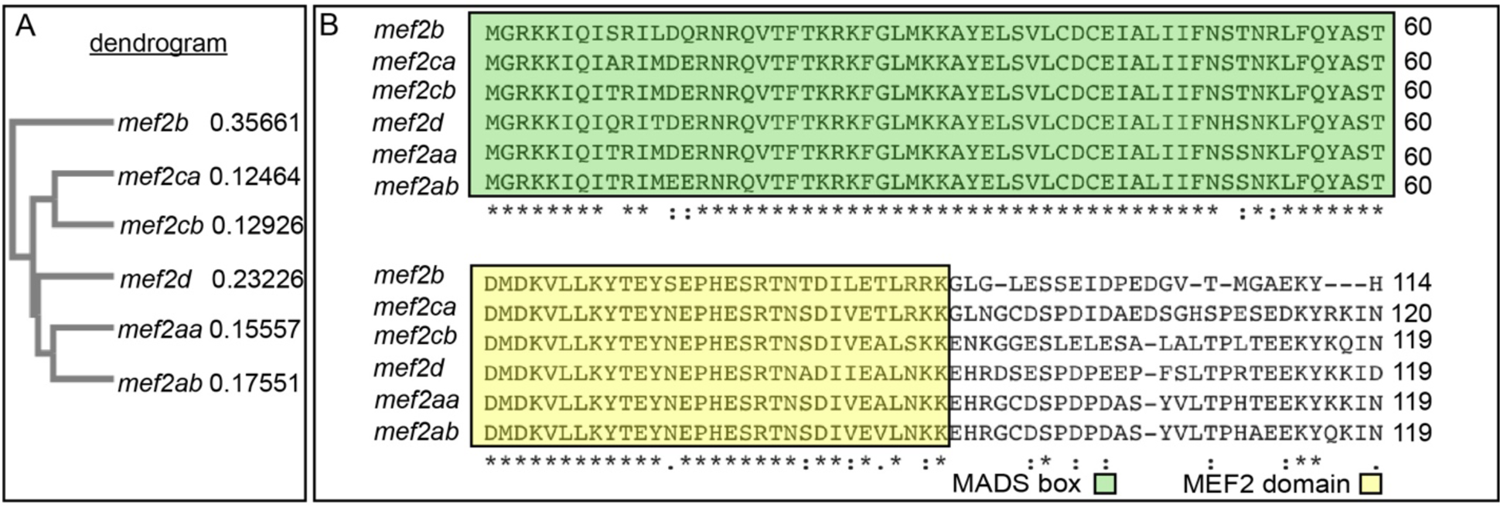
Zebrafish *mef2* paralogs encode highly conserved MADS box and MEF2 domains. (A) Neighbor joining tree generated by Clustal Omega Multiple Sequence Alignment tool depicts the evolutionary relationships between the different zebrafish *mef2* paralogs. The distance values (branch length) are indicated, which represent the evolutionary distance between the individual amino acid sequences and a consensus sequence. (B) *mef2* encoded protein sequence alignment reveals high conservation of MADS box (green) and MEF2 (yellow) domains among all six paralogs. These domains are responsible for DNA binding, dimerization, and cofactor interactions. Transcript IDs used for alignment using the HHalign algorithm are listed in the Materials and Methods.

### Closely related *mef2* paralogs share similar expression dynamics and are downregulated in the high-penetrance strain

To determine the gene expression dynamics of the *mef2* paralogs during craniofacial development, we performed RT-qPCR in wild-type zebrafish heads from 20-30 hpf, stages when *mef2ca* is active^29, 34, 35^ (Fig. 4A). We found that closely-related paralogs share similar gene expression dynamics. *mef2ca* and *mef2cb* have both early and late expression peaks. *mef2aa* and *mef2ab* share an early expression peak. *mef2d* and *mef2b* have a single late and early expression peak, respectively. These findings suggest that there are modules of *mef2* family gene expression in the zebrafish head during the period when the craniofacial skeleton is developing, and that shared regulatory sequences might control expression of different *mef2* gene duplicates.

**Figure 4.**
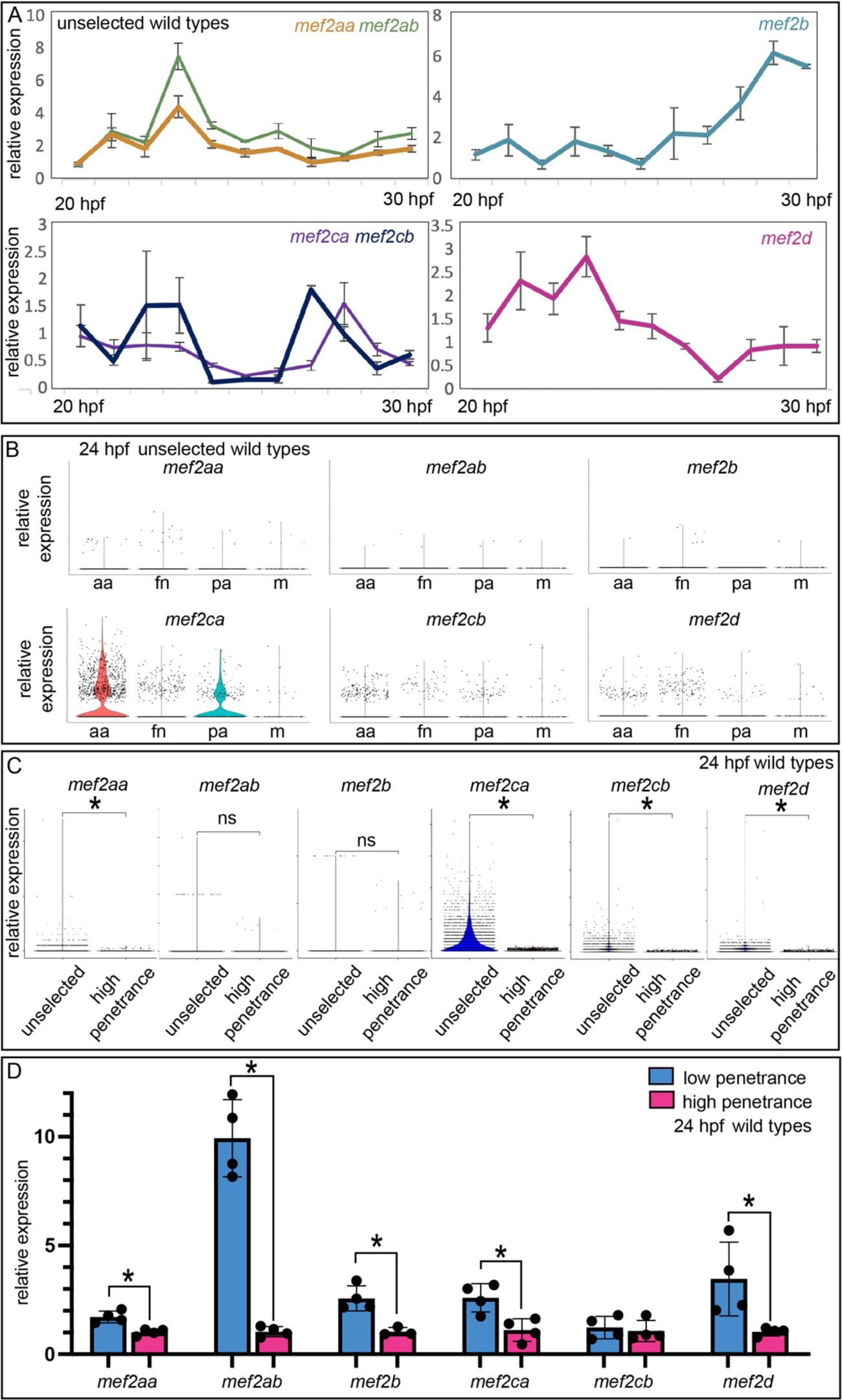
Wild type *mef2* paralog expression dynamics and strain-specific expression levels. (A) Wild-type head expression of all *mef2* paralogs was quantified by RT-qPCR at one hour intervals from 20 to 30 hpf. Expression of each paralog was normalized to *rps18*. Error bars are standard deviation. (B) Expression of each paralog in cranial neural crest cells at 24 hpf was determined by single cell RNA-sequencing on sorted cells. Seurat-based clustering subdivided the cells into four populations as described^41^: anterior arches (aa), frontonasal (fn), posterior arches (pa), and a satellite population containing melanocyte lineage cells (m). (C) Cranial neural crest cell expression of each *mef2* paralog at 24 hpf was quantified by single cell RNA-sequencing. Wild-type animals from an unselected strain were compared to wild-type animals from the high-penetrance selectively bred strain. The scale was adjusted for the unselected strain expression. Asterisks indicate significant differences between strains at p<0.05 by Wilcoxon rank sum test. Differences that were not significantly different (ns) are indicated. (D) Wild-type head expression of all *mef2ca* paralogs was quantified by RT-qPCR at 24 hpf to compare paralog expression levels between the low-penetrance and high-penetrance strains. Expression of each paralog was normalized to *rps18.* Asterisks indicate significant differences at p<0.05. Error bars are standard deviation.

To focus our analysis on just the cells giving rise to the craniofacial skeleton, rather than whole heads, we used single cell RNA-sequencing to examine *mef2* paralog expression in isolated wild-type cranial neural crest cells^41^. Not surprisingly, we found that *mef2ca* has the strongest expression of all the paralogs across different populations of cranial neural crest cells, and that expression is strongest in the anterior arches (Fig. 4B). Other paralogs are more weakly expressed in these cells.

When we demonstrated that paralog transcriptional adaptation does not account for the phenotypic differences between the low- and high-penetrance strains^34^, we did not examine if selective breeding changed paralog expression between strains in *mef2ca* wild types. To examine this possibility, we used single cell RNA-sequencing to compare *mef2* gene expression in cranial neural crest cells between unselected wild types and high-penetrance wild types (Fig. 4C). Strikingly, we found that *mef2aa*, *mef2ca*, *mef2cb* and *mef2d* were all significantly downregulated in the high-penetrance line compared with the unselected strain. These findings suggest that selecting for high penetrance of *mef2ca*-associated phenotypes led to decreased expression of *mef2ca* paralogs. We next used RT-qPCR to compare *mef2* paralog expression between high- and low-penetrance wild types. We found that *mef2aa*, *mef2ab*, *mef2b, mef2ca,* and *mef2d* were all significantly downregulated in high-penetrance heads compared with low-penetrance wild-type heads (Fig. 4D). These findings strongly suggest that one outcome of selective breeding for low- and high-*mef2ca* phenotype penetrance is generally increased and decreased expression, respectively, of the *mef2* paralogs. We do not see overall increases in transcription in the low-penetrance strain compared with the high penetrance strain; housekeeping genes are not significantly upregulated in the low-penetrance strain (Fig. S3). *mef2* paralog downregulation in the high-penetrance strain may explain why *mef2ca*-associated phenotypes occasionally appear in wild types from this strain. We hypothesize that these paralog expression changes underlie the differences in severity and variation between strains.

### *mef2cb* buffers against *mef2ca* loss in the low-penetrance strain

To directly test if upregulated paralog expression buffers the low-penetrance strain against *mef2ca* loss, we systematically removed functional copies of *mef2ca* paralogs from this strain. *mef2cb* is most closely related to *mef2ca*, is the second highest expressed in cranial neural crest cells, and is significantly downregulated in the high-penetrance strain (Figs. 3, 4). We confirmed that *mef2cb* homozygous mutants do not have an overt craniofacial phenotype and are homozygous viable^29^ (Fig. 5A, B, C), indicating that *mef2cb* does not function in zebrafish craniofacial development in an otherwise wild-type background.

**Figure 5.**
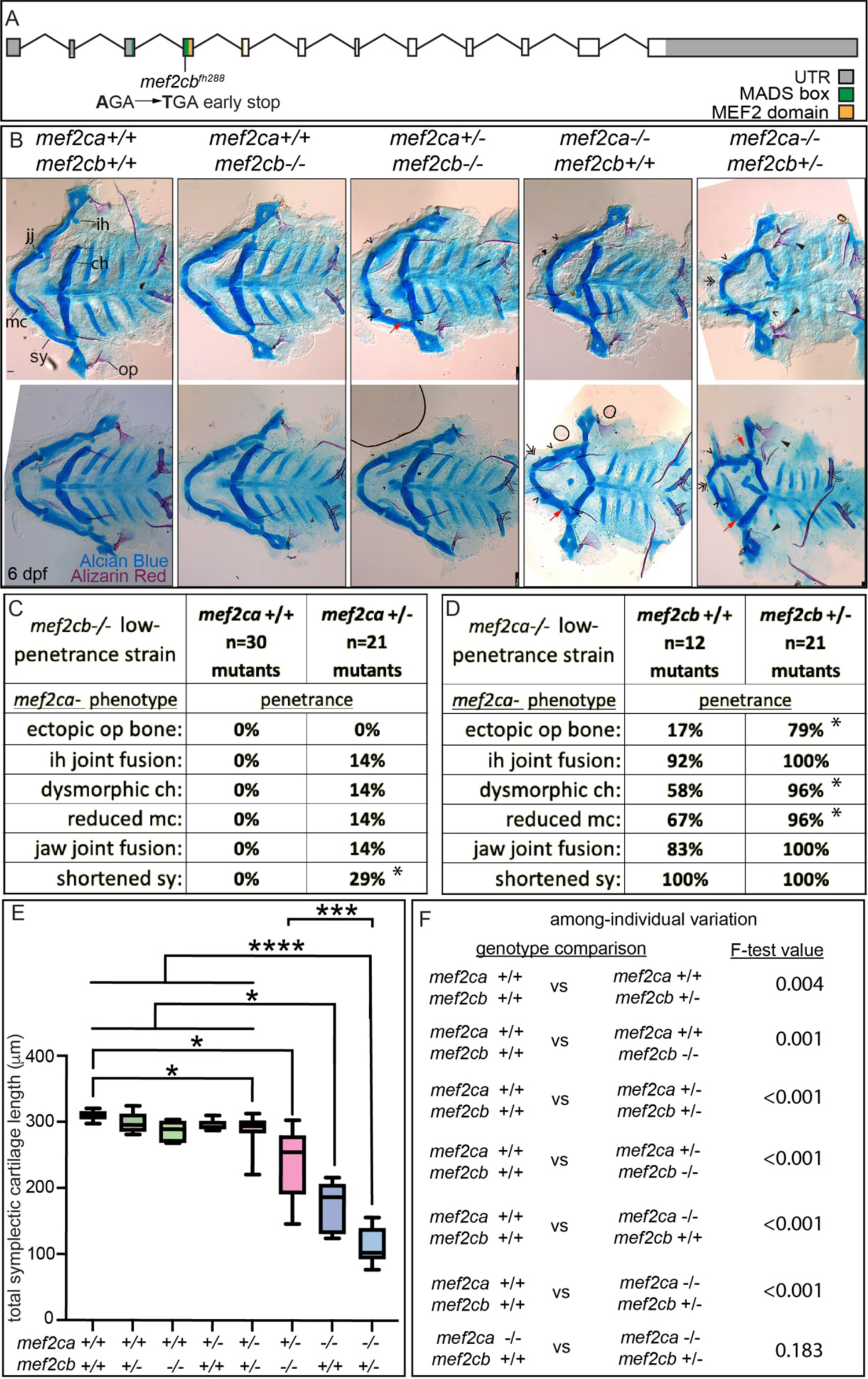
*mef2cb* function buffers against *mef2ca* loss in the-low penetrance strain. (A) Schematic of *mef2cb* exonic structure, mutant allele used in this study and regions encoding proposed functional domains are annotated. (B) Zebrafish heterozygous for both *mef2ca^b1086^* and *mef2cb* were pairwise intercrossed. Six dpf larvae were stained with Alcian Blue and Alizarin Red to label cartilage and bone. Stained larvae were genotyped, flat mounted, and imaged. The following craniofacial skeletal elements are indicated in a wild-type individual: opercle bone (op), branchiostegal ray (br), Meckel’s (mc), ceratohyal (ch), symplectic (sy) cartilages, interhyal (ih) and jaw (jj) joints. Indicated phenotypes associated with *mef2ca* mutants include: ectopic bone (arrowheads), interhyal and jaw-joint fusions (^), dysmorphic ch (arrows), reduced mc (double arrowhead), and a shortened sy (red arrows). Scale bar: 50μm (C, D) The penetrance of *mef2ca* mutant-associated phenotypes observed in 6 dpf larvae are indicated. Asterisk indicates significant difference in penetrance between the indicated genotypes by Fishers exact test. (E) Symplectic cartilage length was measured from 6 dpf larvae from the indicated genotypes. Asterisks indicate significant differences in symplectic length. The p-values from a Dunnet’s T3 test are indicated (* ≤ 0.05, *** ≤ 0.001 **** ≤ 0.0001) (F) Table listing F-test values for significant differences in variation between genotypes. For box and whisker plots, the box extends from the 25th to 75th percentiles. The line in the middle of the box is plotted at the median and the bars are minimum and maximum values. N’s for all analyses are indicated in C and D.

However, when we removed one functional copy of *mef2ca* from *mef2cb* homozygous mutants *mef2ca* mutant-associated phenotypes developed (Fig. 5B, C), phenocopying the high-penetrance strain^34^. We find further evidence for phenocopy; *mef2ca* homozygous mutant phenotype penetrance increased when we removed one copy of *mef2cb* (Fig. 5B, D). Removing both copies of *mef2cb* from *mef2ca* homozygous mutants produces severe, nonspecific defects which make larvae impossible to meaningfully study^29^; although their craniofacial skeletons are severely affected (Fig. S2). Measuring the symplectic cartilage length further demonstrates that when *mef2cb* is fully functional, development is buffered against partial loss of *mef2ca;* there is no difference in symplectic length between wild types and *mef2ca* heterozygotes (wild type vs. *mef2ca^+/-^*;*mef2cb^+/+^*). In contrast, when *mef2cb* is disabled development is sensitive to partial loss of *mef2ca* (wild type vs. *mef2ca^+/-^*;*mef2cb^-/-^*) (Fig. 5E). Removing copies of *mef2cb* from *mef2ca* wild types does not significantly change symplectic cartilage length, but does significantly increase symplectic cartilage variation (Fig. 5F). Thus, even in the *mef2ca* wild-type context, this paralog buffers against phenotypic variation. We conclude that *mef2cb* buffers against *mef2ca-*associated phenotype severity, and among-individual variation, but not within-individual variation (Fig. 5D, E, S3). However, our gene expression study in high- and low-penetrance strains indicated that several *mef2* paralogs are differentially expressed between strains. Therefore, we examined how other paralogs might also buffer against *mef2ca* loss in the low-penetrance strain.

### *mef2d* buffers against *mef2ca* homozygous mutant severity but not variability

We detected *mef2d* transcripts in the anterior arch population of 24 hpf wild-type cranial neural crest cells, and *mef2d* expression is significantly decreased in the high-penetrance strain (Fig. 4C, D). To explore the function of this paralog, we generated a *mef2d* mutant allele in the low-penetrance strain (Fig. 6A). Homozygous *mef2d* mutants did not develop any overt skeletal phenotypes in an otherwise wild-type background (Fig. 6B), and intercrosses between heterozygotes produced homozygous mutant adults at the expected Mendelian frequency, indicating that *mef2d* is not required for viability in laboratory conditions. Similarly, *Mef2d* mutant mice are viable and display no overt phenotypic abnormalities^42, 43^. To test our hypothesis that reduced levels of *mef2d* expression is associated with selective breeding for high *mef2ca* penetrance contributes to severity, we examined offspring from *mef2ca;mef2d* double heterozygous parents (Fig. 6B). When we examined the penetrance of all *mef2ca*-associated mutant skeletal phenotypes in *mef2ca* homozygous mutants, we found that the frequency of ventral cartilage defects was significantly increased by removing a single functional copy of *mef2d,* and further increased in double homozygous mutants (Fig. 6C). The penetrance of other *mef2ca*-associated phenotypes was not significantly changed when *mef2d* function was removed. While symplectic length was not affected in *mef2d* single mutants, we found that removing both copies of *mef2d* from *mef2ca* homozygous mutants significantly shortened symplectic cartilage length. We do not observe significant changes in among-individual variation when we remove *mef2d* function from *mef2ca* homozygotes, but *mef2ca* heterozygotes are more variable when *mef2d* is disabled (Fig. 6E).

**Figure 6.**
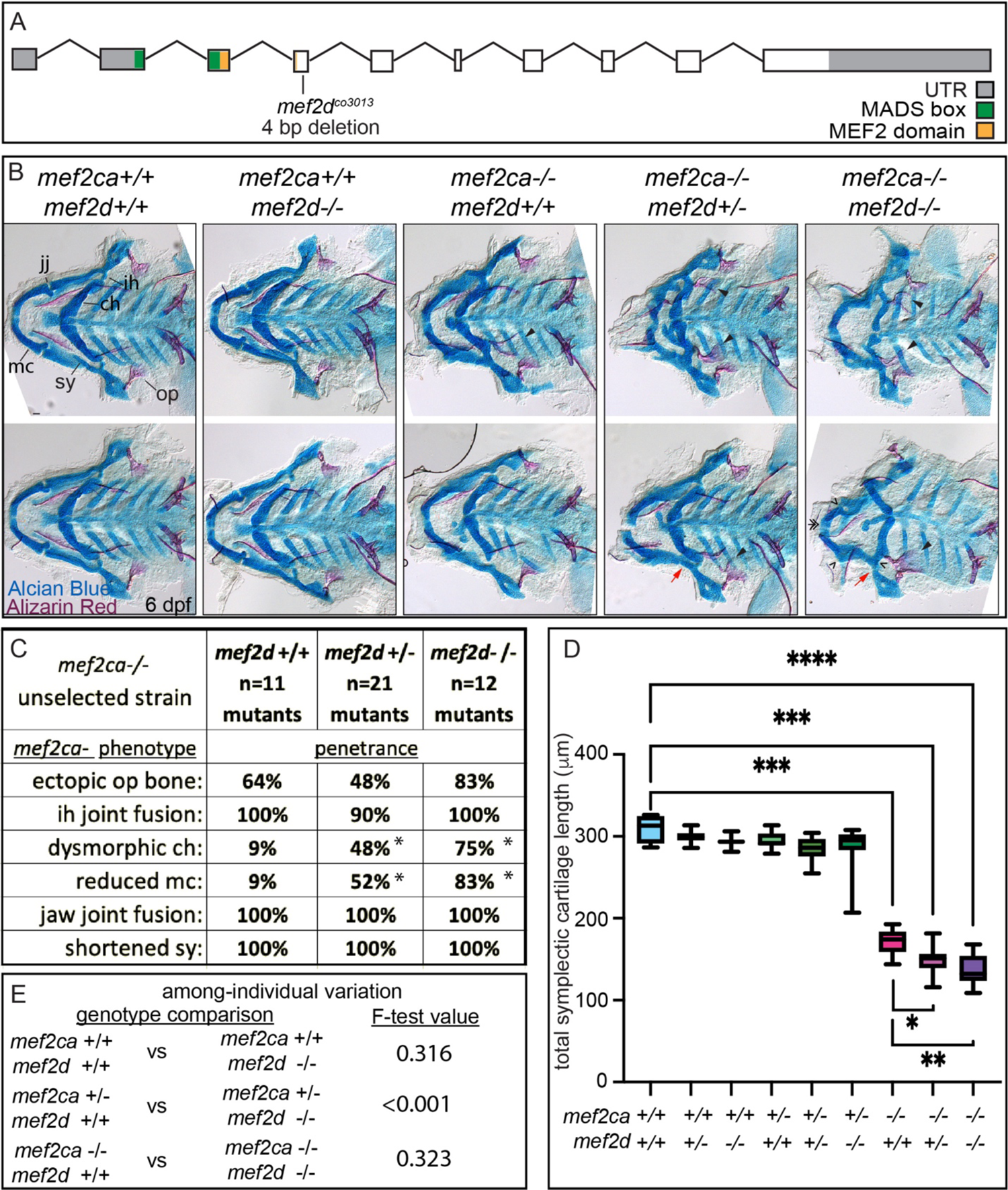
*mef2d* function buffers against *mef2ca* loss in the-low penetrance strain. (A) Schematic of *mef2d* exonic structure, mutant allele used in this study, and regions encoding proposed functional domains are annotated. (B) Zebrafish heterozygous for both *mef2ca ^b1086^* and *mef2d* were pairwise intercrossed. Six dpf larvae were stained with Alcian Blue and Alizarin Red to label cartilage and bone. Stained larvae were genotyped, flat mounted, and imaged. The following craniofacial skeletal elements are indicated in a wild-type individual: opercle bone (op), branchiostegal ray (br), Meckel’s (mc), ceratohyal (ch), symplectic (sy) cartilages, interhyal (ih) and jaw (jj) joints. Indicated phenotypes associated with *mef2ca* mutants include: ectopic bone (arrowheads), interhyal and jaw-joint fusions (^), dysmorphic ch (arrows), reduced mc (double arrowhead), and a shortened sy (red arrows). Scale bar: 50μm (C) The penetrance of *mef2ca* mutant-associated phenotypes observed in 6 dpf larvae are indicated. Asterisk denotes significant difference in penetrance between the indicated genotypes by Fishers exact test. (D) Symplectic cartilage length was measured from 6 dpf larvae from the indicated genotypes, asterisk indicates significant differences in symplectic length (* ≤ 0.05, ** ≤ 0.01 *** ≤ 0.001). (E) Table listing F-test values testing for significant differences in variation between genotypes. For box and whisker plots, the box extends from the 25th to 75th percentiles. The line in the middle of the box is plotted at the median and the bars are minimum and maximum values. N’s for all analyses are indicated in C.

### *mef2b* buffers against *mef2ca* homozygous mutant variation, but not severity

*mef2b* is the most divergent *mef2ca* paralog, and is only minimally expressed in cranial neural crest cells at 24 hpf (Figs. 3,4). However, we did observe significantly higher *mef2b* expression in the low-penetrance strain compared with the high-penetrance strain (Fig. 4D). We generated a *mef2b* mutant allele in the low-penetrance strain (Fig. 7A). Homozygous *mef2b* mutants did not exhibit any overt skeletal phenotypes (Fig. 7B) and intercrosses between animals heterozygous for this allele produced homozygous mutant adults at the expected Mendelian frequency. When we removed functional copies of *mef2b* from *mef2ca* homozygous mutants, we did not observe any significant changes in *mef2ca* mutant-associated phenotype penetrance (Fig. 7C), nor did we detect further reductions in symplectic cartilage length when functional copies of *mef2b* were removed from *mef2ca* homozygous mutants (Fig. 7D). These findings indicate that *mef2b* function does not affect *mef2ca* mutant penetrance or severity. However, we do observe a modest increase in among-individual variation in *mef2ca* homozygous mutants when *mef2b* is disabled (Fig. 7E).

**Figure 7.**
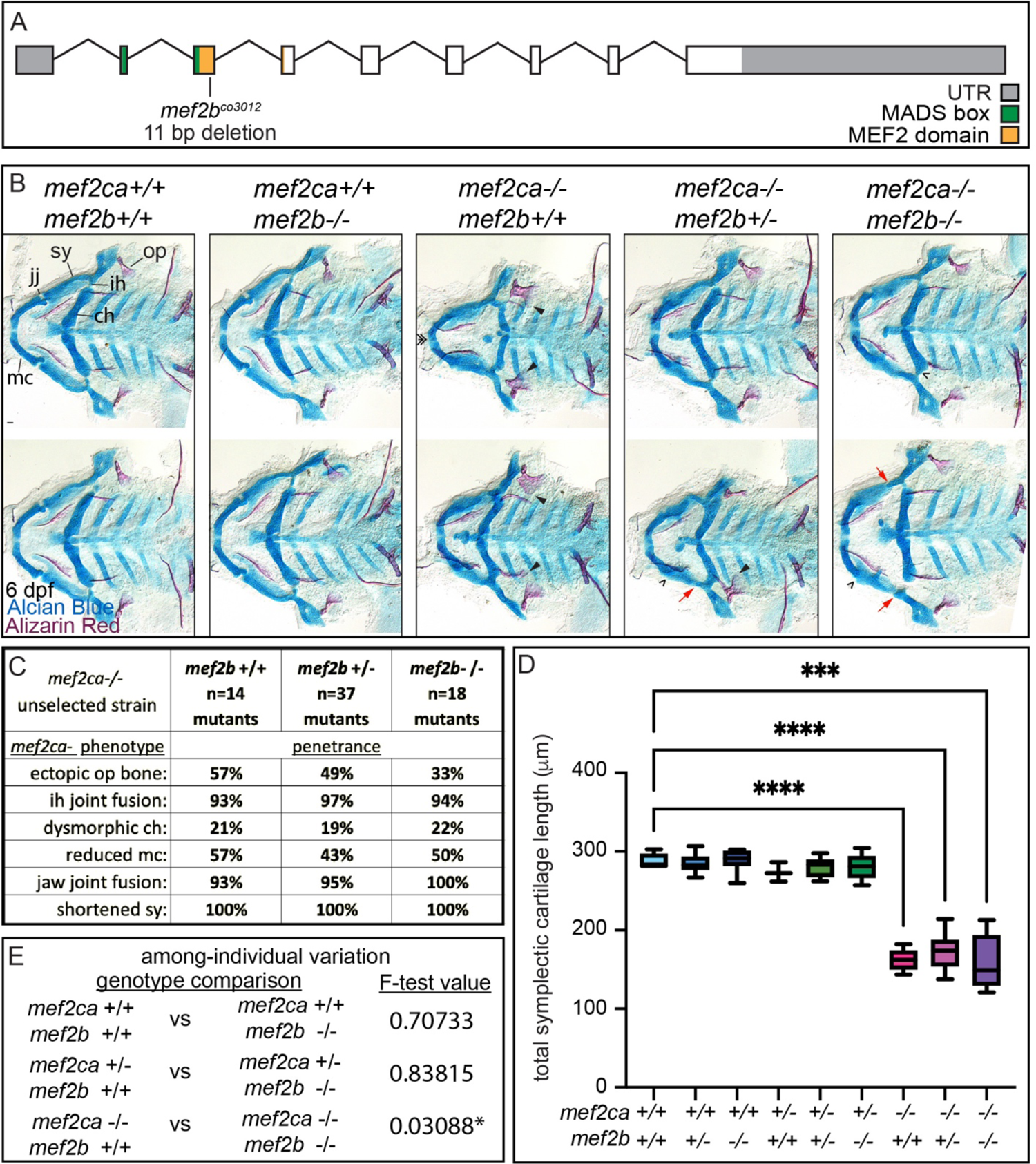
*mef2b* function buffers against *mef2ca* loss in the-low penetrance strain. (A) Schematic of *mef2b* exonic structure, mutant allele used in this study and regions encoding proposed functional domains are annotated. (B) Zebrafish heterozygous for both *mef2ca ^b1086^* and *mef2b* were pairwise intercrossed. Six dpf larvae were stained with Alcian Blue and Alizarin Red to label cartilage and bone. Stained larvae were genotyped, flat mounted, and imaged. The following craniofacial skeletal elements are indicated in a wild-type individual: opercle bone (op), branchiostegal ray (br), Meckel’s (mc), ceratohyal (ch), symplectic (sy) cartilages, interhyal (ih) and jaw (jj) joints. Indicated phenotypes associated with *mef2ca* mutants include: ectopic bone (arrowheads), interhyal and jaw-joint fusions (^), dysmorphic ch (arrows), reduced mc (double arrowhead), and a shortened sy (red arrows). Scale bar: 50μm (C) The penetrance of *mef2ca* mutant-associated phenotypes observed in 6 dpf larvae are indicated. (D) Symplectic cartilage length was measured from 6 dpf larvae from the indicated genotypes, asterisk indicates significant differences in symplectic length (*** ≤ 0.001 **** ≤ 0.0001). (E) Table listing F-test values testing for significant differences in variation between genotypes. For box and whisker plots, the box extends from the 25th to 75th percentiles. The line in the middle of the box is plotted at the median and the bars are minimum and maximum values. N’s for all analyses are indicated in C.

### *mef2aa* buffers against *mef2ca* heterozygous phenotypes, but not *mef2ca* homozygous mutant phenotypes

*mef2aa* is minimally expressed in cranial neural crest cells (Figs. 3, 4). However, *mef2aa* expression differs between strains (Fig. 4C, D). We developed a *mef2aa* mutant (Fig. 8A) that does not develop any overt skeletal phenotypes when homozygous (Fig. 8B). When we removed *mef2aa* from *mef2ca* homozygous mutants, we did not observe any significant changes in *mef2ca*-associated phenotype penetrance (Fig. 8C), symplectic cartilage length (Fig. 8D), or variation (Fig. 8E). However, when we removed one functional copy of *mef2ca* from *mef2aa* homozygous mutants, *mef2ca*-associated phenotypes developed with low frequency (Fig. 8B). Specifically, some animals developed nubbins and shortened symplectic cartilages. Both of these phenotypes are never seen in unselected *mef2ca* heterozygotes, but do develop in *mef2ca* heterozygotes from the high-penetrance strain^34^. Therefore, disabling *mef2aa* partially phenocopies the high-penetrance strain.

**Figure 8.**
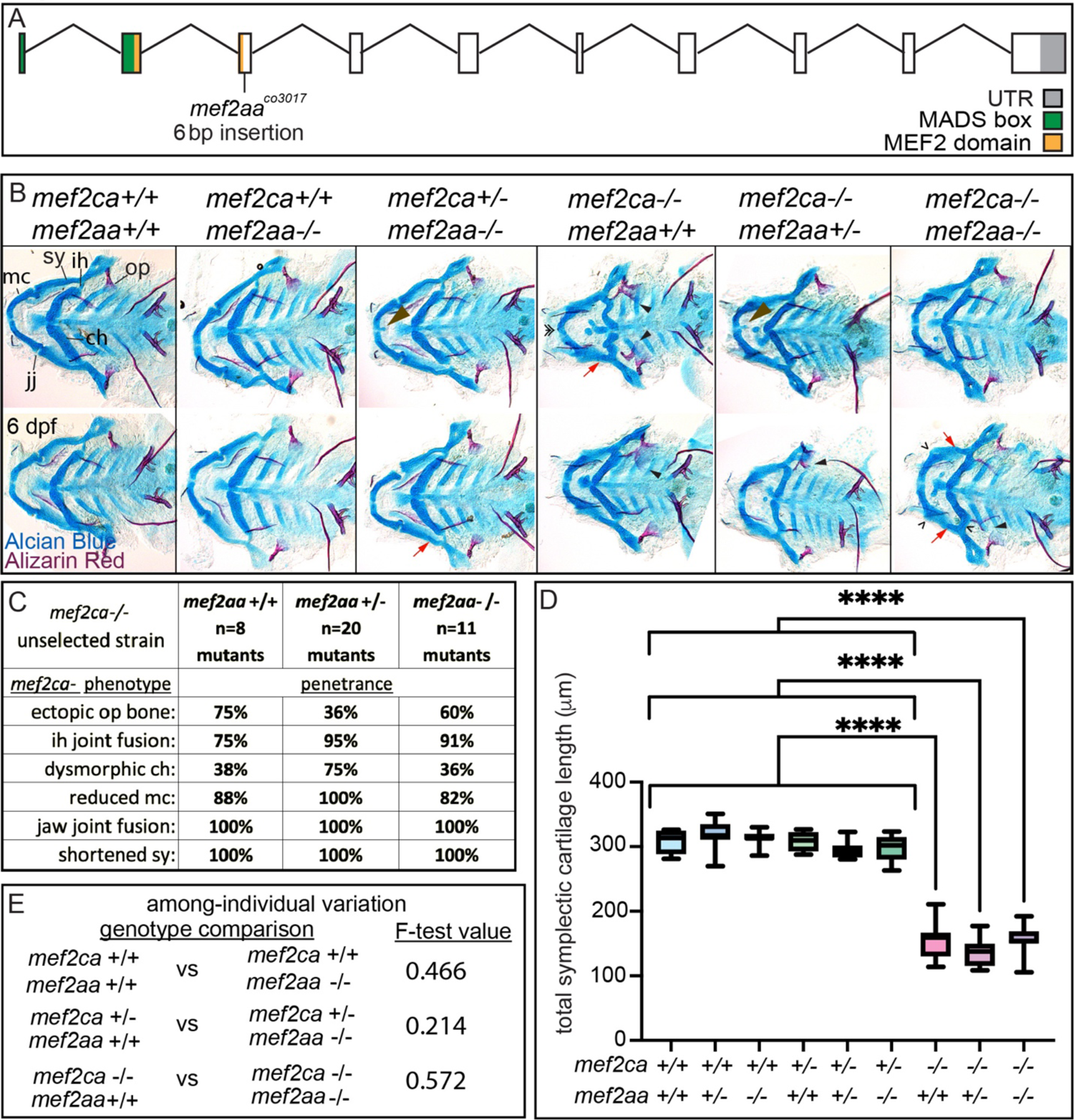
*mef2aa* function buffers against *mef2ca* partial loss in the-low penetrance strain. (A) Schematic of *mef2aa* exonic structure, mutant allele used in this study, and regions encoding proposed functional domains are annotated. (B) Zebrafish heterozygous for both *mef2ca ^b1086^* and *mef2aa* were pairwise intercrossed. Six dpf larvae were stained with Alcian Blue and Alizarin Red to label cartilage and bone. Stained larvae were genotyped, flat mounted, and imaged. The following craniofacial skeletal elements are indicated in a wild type: opercle bone (op), branchiostegal ray (br), Meckel’s (mc), ceratohyal (ch), symplectic (sy) cartilages, interhyal (ih) and jaw (jj) joints. Indicated phenotypes associated with *mef2ca* mutants include: cartilage nubbin fused to the mc symphysis (brown arrowhead), ectopic bone (black arrowheads), interhyal and jaw-joint fusions (^), dysmorphic ch (arrows), reduced mc (double arrowhead), and a shortened sy (red arrows). Scale bar: 50μm (C) The penetrance of *mef2ca* mutant-associated phenotypes observed in 6 dpf larvae are indicated. (D) Symplectic cartilage length was measured from 6 dpf larvae from the indicated genotypes, asterisk indicates significant differences in symplectic length (**** ≤ 0.0001). (E) Table listing F-test values testing for significant differences in variation between genotypes. For box and whisker plots, the box extends from the 25th to 75th percentiles. The line in the middle of the box is plotted at the median and the bars are minimum and maximum values. N’s for all analyses are indicated in C.

## DISCUSSION

### Vestigial paralog expression dynamics may buffer development

Vertebrate *mef2* functions downstream of Endothelin signaling in the developing craniofacial skeleton^35, 44^. The endothelin pathway was subfunctionalized following whole genome duplications in vertebrates^45^. Thus, *mef2* genes may have been subfunctionalized following genome duplications. Comparing mice and zebrafish further supports *mef2* subfunctionalization. In mice, *Mef2c* is required for both heart and craniofacial development^44, 46^. In zebrafish, both co-orthologs (*mef2ca* and *mef2cb*) function redundantly in the heart^28^, while craniofacial function has been subfunctionalized to just *mef2ca* (Fig. 9). However, it is possible that the ancestral craniofacial function and expression pattern of *mef2cb* is partially retained following subfunctionalization, even though it is no longer required for this function. We propose that while duplicated genes can evolve new expression domains and functions, vestiges of their original expression pattern remain and can buffer against loss of another paralog. Consistently, the *mef2* paralog mutants don’t have an overt skeletal phenotype except for enhancing specific *mef2ca* mutant phenotypes. In our system, selective breeding partially resurrected paralog ancestral expression and restored near full redundancy in the low-penetrance strain (Fig. 9). Our experiments support a model where we selected upon alleles that either amplify or dampen existing vestigial expression depending on the direction of selection. This model predicts that pre-existing genomic variation in *mef2* paralog enhancers, which is cryptic in wild types, underlies the phenotypic variation we observe, and is what we selected upon in *mef2ca* mutants.

**Figure 9.**
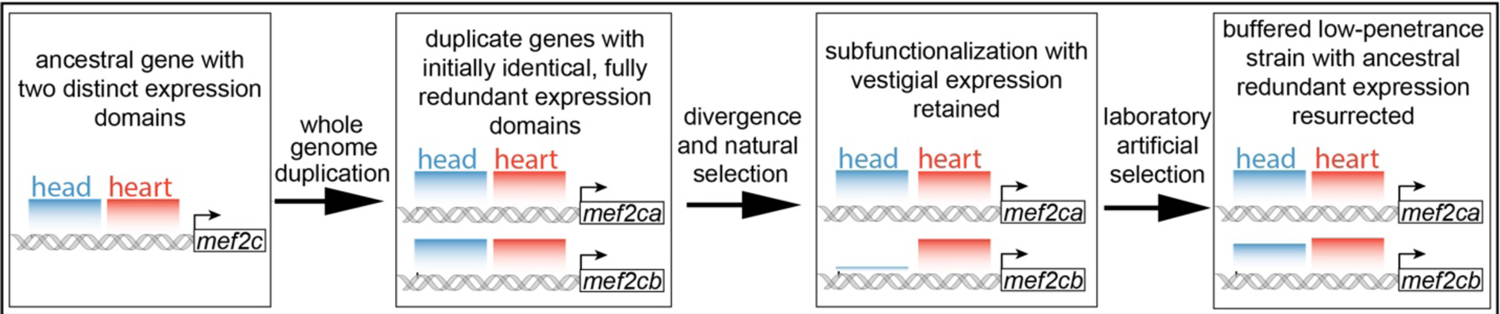
Model for *mef2* gene duplicate evolution and resurrection of ancestral expression via selective breeding. In our model, an ancestral *mef2* gene existed with distinct regulatory regions. In this example, ancestral *mef2c* had enhancers driving expression in the head (blue) and the heart (red). Following whole genome duplication, initially the regulatory and coding sequences would be identical. Through divergence and natural selection, regulatory sequences would acquire mutations that dampen, but do not eliminate, some expression domains. In this example *mef2cb* expression is dampened in the head. These gene expression changes result in subfunctionalization, but with vestigial retention of the original expression. *mef2cb* is no longer required for craniofacial development, yet traces of the original craniofacial expression remain. The amount of vestigial expression is variable among individuals, resulting in variable buffering against *mef2ca* mutant phenotypes across a population. Our selective breeding selected on this variable, expression resulting in increased paralog expression in the low-penetrance strain. Thus, *mef2* paralog expression in the low-penetrance strain resembles the ancestral condition following whole genome duplication, when both copies’ expression profiles were highly similar and the genes, were redundant for craniofacial development.

### Decoupling buffering mechanisms

We found that paralogs modularly buffer the *mef2ca* homozygous mutants. For example, disabling *mef2cb* affects most phenotypes, while *mef2d* mutations primarily affect penetrance of ventral cartilages, and only variation is affected when *mef2b* function is removed. Further, expression of all the paralogs was affected by selection, indicating that they all contribute to buffering.

However, we reported that the rapid response to selection indicates that relatively few heritable factors contribute to penetrance variation^27, 34^. We propose that a few, degenerate enhancers shared by the paralogs exist and by selecting for penetrance all paralogs were up or down regulated via these shared regulatory regions. We found some evidence of shared gene regulation in our experiments examining *mef2* gene expression dynamics. Closely related paralog expression dynamics are similar, suggesting shared regulatory sequences. In this model, few enhancers are inherited that control the expression of many paralogs.

Determining whether the aspect of the *mef2ca* phenotype buffered by each paralog is related to their wild type expression pattern would expand our understanding of modularity, although *in situ* gene expression protocols were not sensitive enough to detect the low expression of the individual paralogs. Moreover, the quantitative phenotyping in this study was limited to the symplectic cartilage. Quantitative phenotyping of other parts of the developing craniofacial complex^47, 48^ would likely reveal more information about modular buffering of severity and variation by paralogs. For example, penetrance of ceratohyal cartilage defects was affected by *mef2d*, and therefore variation of this structure, which we did not measure, might be affected by loss of *mef2d*.

Our experiments also decouple the mechanisms that buffer among- and within-individual variation. Examining allele severity, strain severity, and the different paralog mutants demonstrate buffering in our system only regulates among-individual variation, not within-individual variation (Fig. S4). These two types of variation are buffered by different mechanisms in our system.

These studies advance our understanding of craniofacial variation. We propose that cryptic variation in vestigial paralog expression is the noise underlying variable craniofacial development. This paralog expression variation may explain how some genetically resilient individuals can overcome a deleterious mutant allele.

## FIGURES AND LEGENDS

**Figure S1.**
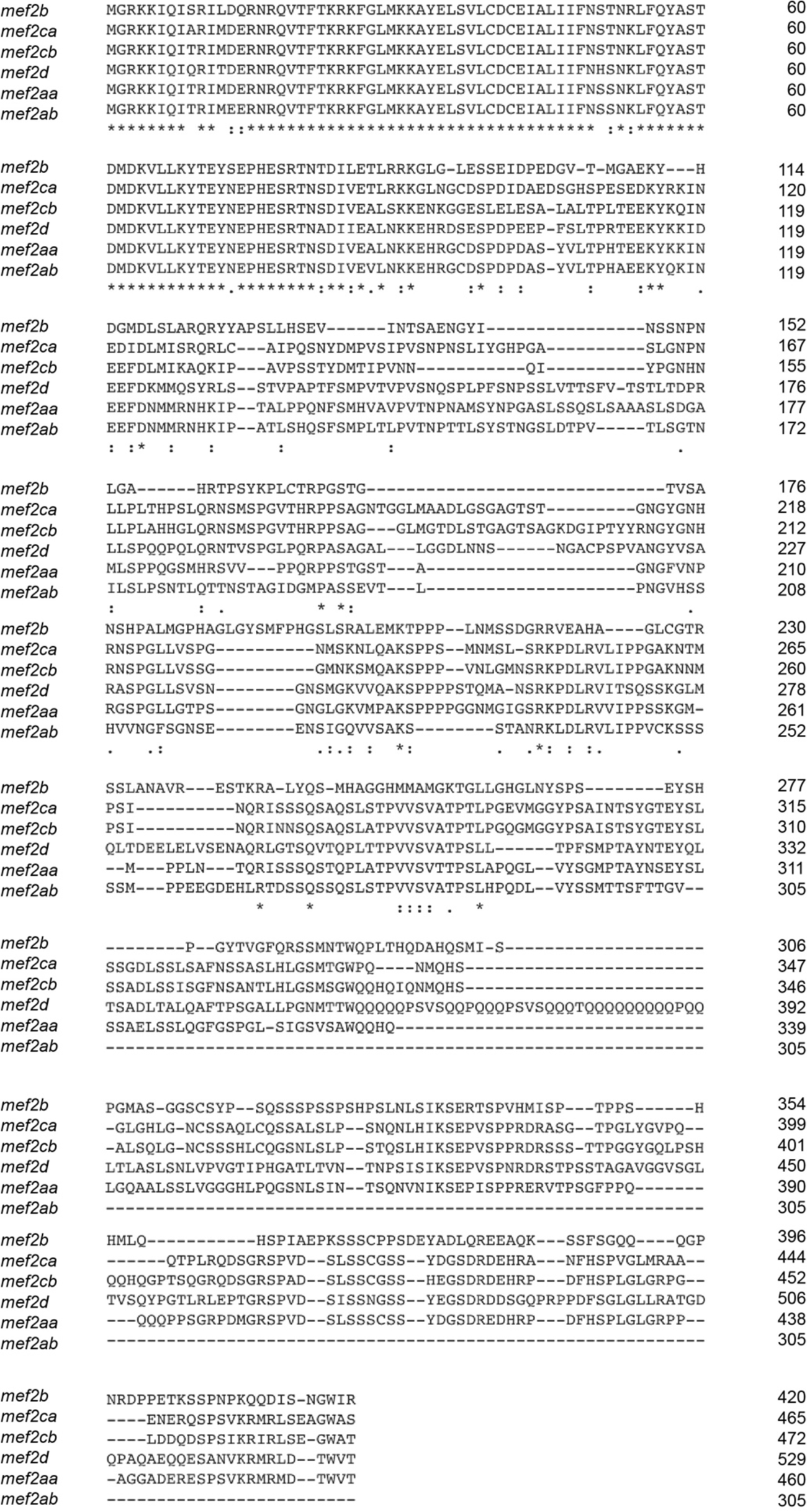
Zebrafish *mef2* paralogs encode highly conserved N-terminal MADS box and MEF2 domains, but divergent C-terminal domains. *mef2* encoded protein sequence alignment reveals high conservation of MADS box and MEF2 domains among all six paralogs. However, the rest of the proteins encoded by these genes are poorly conserved. Transcript IDs used for alignment using the HHalign algorithm are listed in the Materials and Methods.

**Figure S2.**
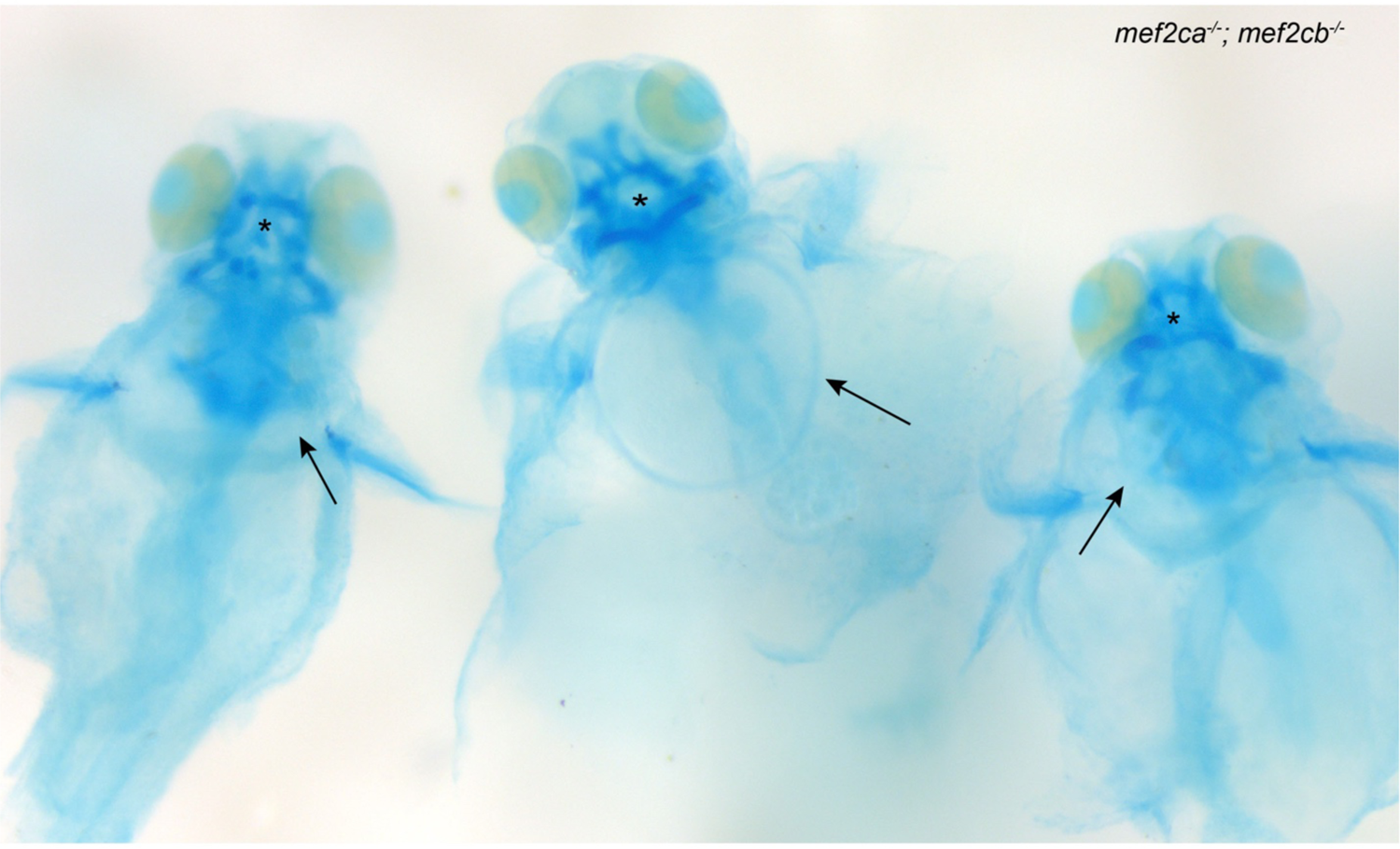
Zebrafish *mef2ca;mef2cb* double homozygous mutants develop nonspecific developmental defects. Zebrafish double heterozygous for *mef2ca^b1086^* and *mef2cb* were intercrossed. Offspring were stained with Alcian Blue and Alizarin Red then genotyped for both genes. In whole-mount images of double homozygous mutants severe edema (arrows), and craniofacial cartilage malformations (asterisks) are observed with 100% penetrance.

**Figure S3.**
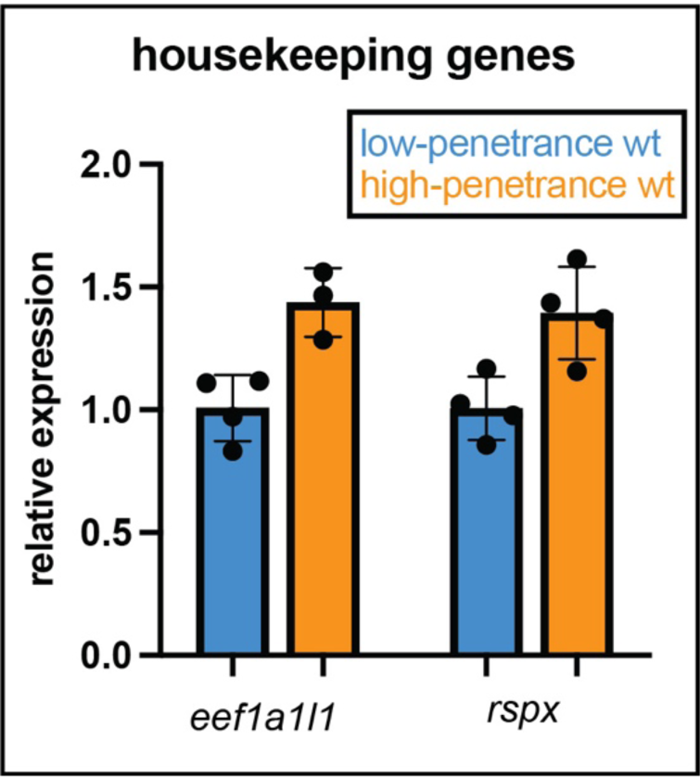
Overall transcription is not increased in the low-penetrance strain. We observed increased expression of many *mef2* paralogs in wildtypes from the low-penetrance strain compared with high-penetrance strain wildtypes. To determine if overall gene expression is generally upregulated in the low-penetrance strain, we quantified two housekeeping genes by RT-qPCR at 24 hpf to compare expression levels between strains. Expression of each gene was normalized to *rps18.* In contrast to the *mef2* paralogs, neither housekeeping gene is significantly upregulated in the low-penetrance strain. Error bars are standard deviation.

**Figure S4.**
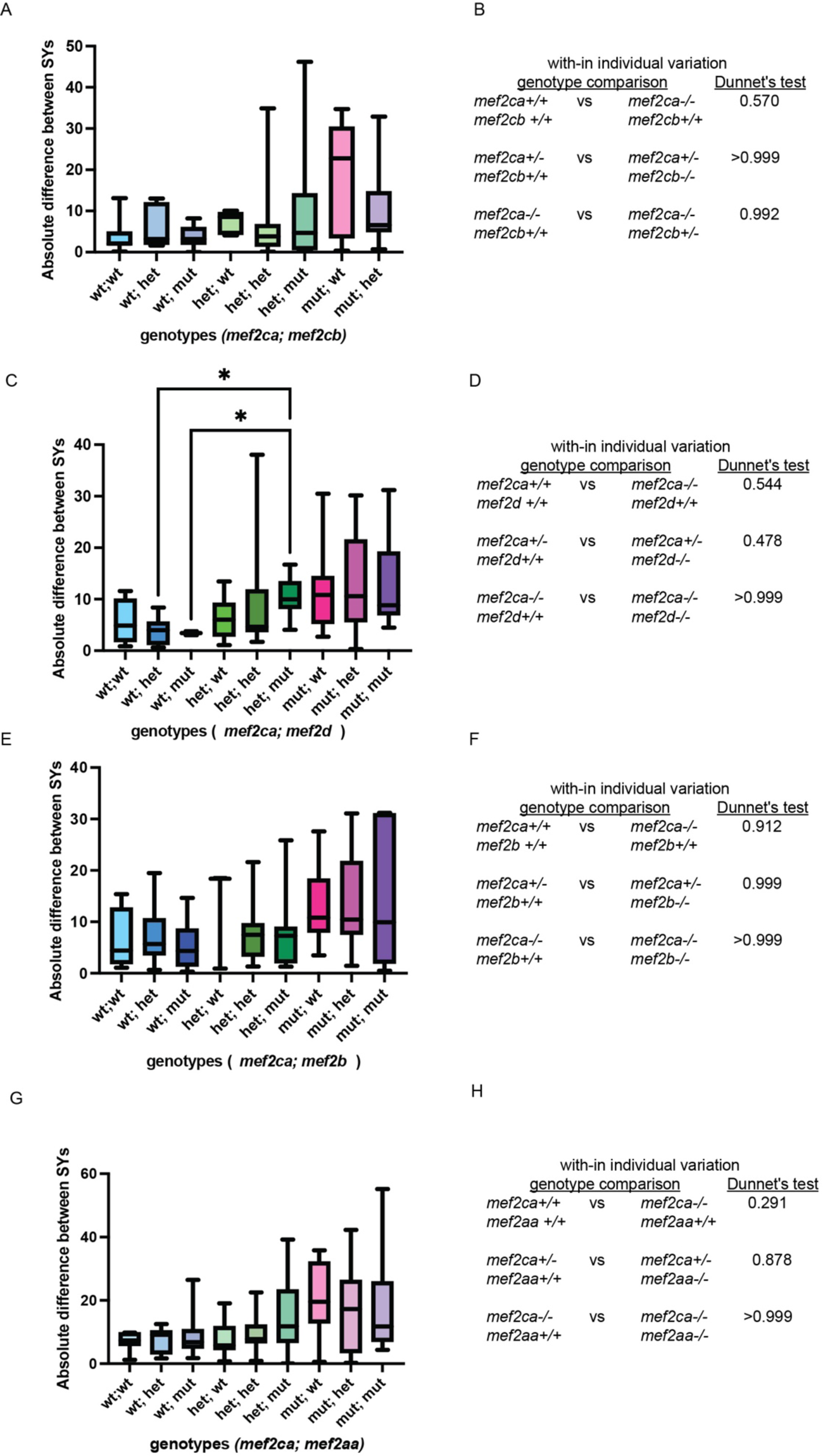
Within-individual variation does not correlate with severity. (A, C, E, G) Symplectic cartilage length on left and right sides of 6 dpf zebrafish stained with Alcian Blue and Alizarin Red was measured to determine fluctuating asymmetry or the absolute difference between left and right for fish with the indicated genotypes. (B, D, F, H) Table listing the Dunnett’s test values for significant differences in fluctuating asymmetry between fish with the indicated genotypes. For box and whisker plots, the box extends from the 25th to 75th percentiles. The line in the middle of the box is plotted at the median and the bars are minimum and maximum values.

## METHODS

### Zebrafish strains and husbandry

All fish were maintained and staged according to established protocols^49,50^. Selective breeding was performed as previously described^27,39^. The *mef2ca^b631^*, *mef2ca^b1086^*, and *mef2cb^fh288^* mutant alleles have been previously described^28,35^ and were maintained by outcrossing to the AB background or to the low-penetrance *mef2ca^b1086^* selectively bred strain.

### CRISPR/Cas9-induced mutant alleles

We generated germline mutant alleles using CRISPR/Cas9 mutagenesis^51^ with modifications as described^41^. Briefly, we designed sgRNAs within or just downstream of the MADS or MEF2 domain. The XbaI-digested pT3TS-nCas9n plasmid (Addgene plasmid #46757) was used as a template to transcribe Cas9 mRNA with the T3 mMESSAGE kit (Invitrogen). We transcribed sgRNAs (see table below) from PCR-generated templates using the MEGAscript T7 Kit (Thermo Fisher Scientific). One cell-stage embryos were injected with a mix of 200 ng/µl Cas9 mRNA and 50 ng/µl of each gene-specific sgRNA. Injected embryos were raised and founders identified by amplifying the genomic region containing the sgRNA site and identifying banding size shifts indicating insertions and/or deletions (see Table xx for primers). All paralog mutants were maintained by backcrossing to the low-penetrance strain for at least three generations. The following sgRNAs were used: *mef2d^co3013^*: 5’-GGACAAATACCGGAAGAGCG-3’ *mef2b^co2012^*: 5’-CACGAGAGCCGCACTAACAC-3’ *mef2aa^co3017^*: 5’-TCATGGACGACCGTTTCGGC-3’ *mef2ca^co3008^*: 5’-GGCTCCAAACTCTATATGGG-3’ and 5’-TCTCCTTCCTCTGTCGTTCC-3’. We generated six independent sgRNAs for *mef2ab* and none of them mutagenized this locus.

### Cartilage and bone staining and imaging

Fixed animals were stained with Alcian Blue and Alizarin Red as described previously^39,52^. Alcian Blue and Alizarin Red-stained 6dpf skeletons were dissected and flat mounted for Nomarski imaging on a Leica DMi8 inverted microscope equipped with a Leica DMC2900 as previously described^53^.

### Phenotype scoring

For penetrance scoring, 6 dpf Alcian Blue and Alizarin Red stained skeletons were genotyped then scored for the proportion of animals with a given genotype that exhibit a particular phenotype. In the interest of strong rigor and reproducibility, phenotypes were scored by three observers blinded to genotype. All three agreed with the number of animals in each phenotypic class, indicating that phenotype penetrance can be reproducibly identified by different observers. For symplectic cartilage measurements, whole mount Alcian Blue, Alizarin Red stained skeletons were imaged under a transmitted light dissecting scope. Images were captured with Zeiss ZEN software. This software was then used to measure the linear distance in microns from the posterior most point of the interhyal cartilage and the distal tip of the symplectic cartilage similar to a previous study^54^. For each individual, the total symplectic length was calculated by summing the measurements from the left and right side. These same measurements were also used to calculate the absolute value of the left minus the right symplectic cartilage lengths to determine developmental instability for each individual. All raw phenotype data are presented in supplementary data tables.

### RT-qPCR

Gene expression studies were performed as previously described^34^. For the time course study from unselected AB, and for comparing low- and high-penetrance wild types, live individual 24 hpf embryos from each strain had their heads removed. Decapitated bodies were genotyped to identify homozygous wild-types. Heads from 5–6 identified homozygous wild types were pooled and total RNA was extracted with TRI Reagent. cDNA was prepared with Superscript III from Invitrogen. qPCR experiments utilized a Real-Time PCR StepOnePlus system from Applied Biosystems and SYBR green. A standard curve was generated from serially diluted (1:2:10) cDNA pools, and primers with a slope of −3.3 +-0.3 were accepted. The relative quantity of target cDNA was calculated using Applied Biosystems StepOne V.2.0 software and the comparative Ct method. After surveying the expression of many housekeeping genes at multiple stages, we determined that *rps18* expression was the most consistent across stages, genotypes, and strains. Target gene expression in all experiments was normalized to *rps18*. Reactions were performed in technical triplicate, and the results represent two to six biological replicates. The following primers were used: *rps18* FW, 5’-CTGAACAGACAGAAGGACATAA-3’ and *rps18* REV 5’-AGCCTCTCCAGATCTTCTC-3’, *mef2ca* FW, 5’-GTCCAGAATCCGAGGACAAATA-3’ and *mef2ca* REV 5’-GAGACAGGCATGTCGTAGTTAG-3’, *mef2cb* FW, 5’-AGTACGCCAGCACAGATA-3’ and *mef2cb* REV 5’-AGCCATTTAGACCCTTCTTTC-3’, *mef2aa* FW, 5’-CCACGAGAGCAGAACCAACTC-3’ and *mef2aa* REV 5’-GTCCATGAGGGGACTGTGAC-3’, *mef2ab* FW, 5’-AACCTCACGAGAGCAGAACC-3’ and *mef2ab* REV 5’-AGGACATATGAGGCGTCTGG-3’, *mef2b* FW, 5’-CCGATATGGACAAAGTGCTG-3’ and *mef2b* REV 5’-CCAATCCCAATCCTTTCCTT-3’, *mef2d* FW, 5’-TTCCAGTATGCCAGCACTGA-3’ and *mef2d* REV 5’-CGAATCACGGTGCTCTTTCT-3’. All qPCR numerical data and statistical analyses are reported in supplementary data table.

### Cell sorting

32 high-penetrance or 60 unselected wild-type *flia:EGFP*;*sox10:mRFP* double transgenic embryos were dissociated into single-cell suspension using cold-active protease from *Bacillus licheniformis*, DNase, EDTA and trituration^41^. Approximately 19,000 or 30,000 live, double-positive cells were sorted into tubes pre-coated with fetal bovine serum with a MoFlo XDP100. Sorted cells at 2.94×10^5^ cells/ml with 85% viability or 2.08×10^6 cells/ml with 70% viability were loaded into the 10x Chromium Controller aiming to capture 10000 cells. Using the NovaSEQ6000, 2×150 paired-end sequencing Chromium 10x Genomics libraries were sequenced at 100K reads per cell using 3prime NexGem V3.1(Dual Index) sequencing.

### scRNA-seq analysis

Recovered sequencing reads were mapped to a custom Cell Ranger reference based on the GRCz11/danRer11 genome assembly using the Cell Ranger pipeline (10X Genomics). The 10X Genomics Cell Ranger pipeline removed empty droplets from the dataset based on the distribution of cell barcodes. Additional analysis was performed in R using the Seurat package^55^. Seurat objects containing two different wildtype populations high-penetrance and unselected were independently quality filtered. Quality filtering of these data excluded cells with no genes detectable (>0 unique molecular identifiers (UMIs) per gene). Each gene in the dataset was required to be present in a minimum of five cells. Regarding the capture of dead or dying cells, a minimum presence of 250 genes (at <2.5% of mitochondrial UMIs detected) were required. After quality filtering, 11257 cells (3329 high-penetrance *mef2ca^+/+^* cells and 7928 unselected *mef2ca^+/+^* cells) remained for analysis with an average of 3810.887 genes per cell. After merging each of the wildtype datasets into a single Seurat object, Seurat’s ‘VlnPlot’ function was used to generate *mef2* paralog plots indicating the distribution of cells expressing each paralog across each of the four total, cell-type clusters. One wild-type dataset analyzed in this study was previously published by us^41^. The raw, feature-barcode matrix for this dataset can be accessed from the GEO database (accession number GSE163826). The second dataset was generated for this study and has been uploaded to GEO (accession number not yet available, but will be provided upon publication).

### Genotyping Assays

*mef2cab1086* was genotyped by KASP as previously described^39^. PCR-based genotyping assays are as follows:

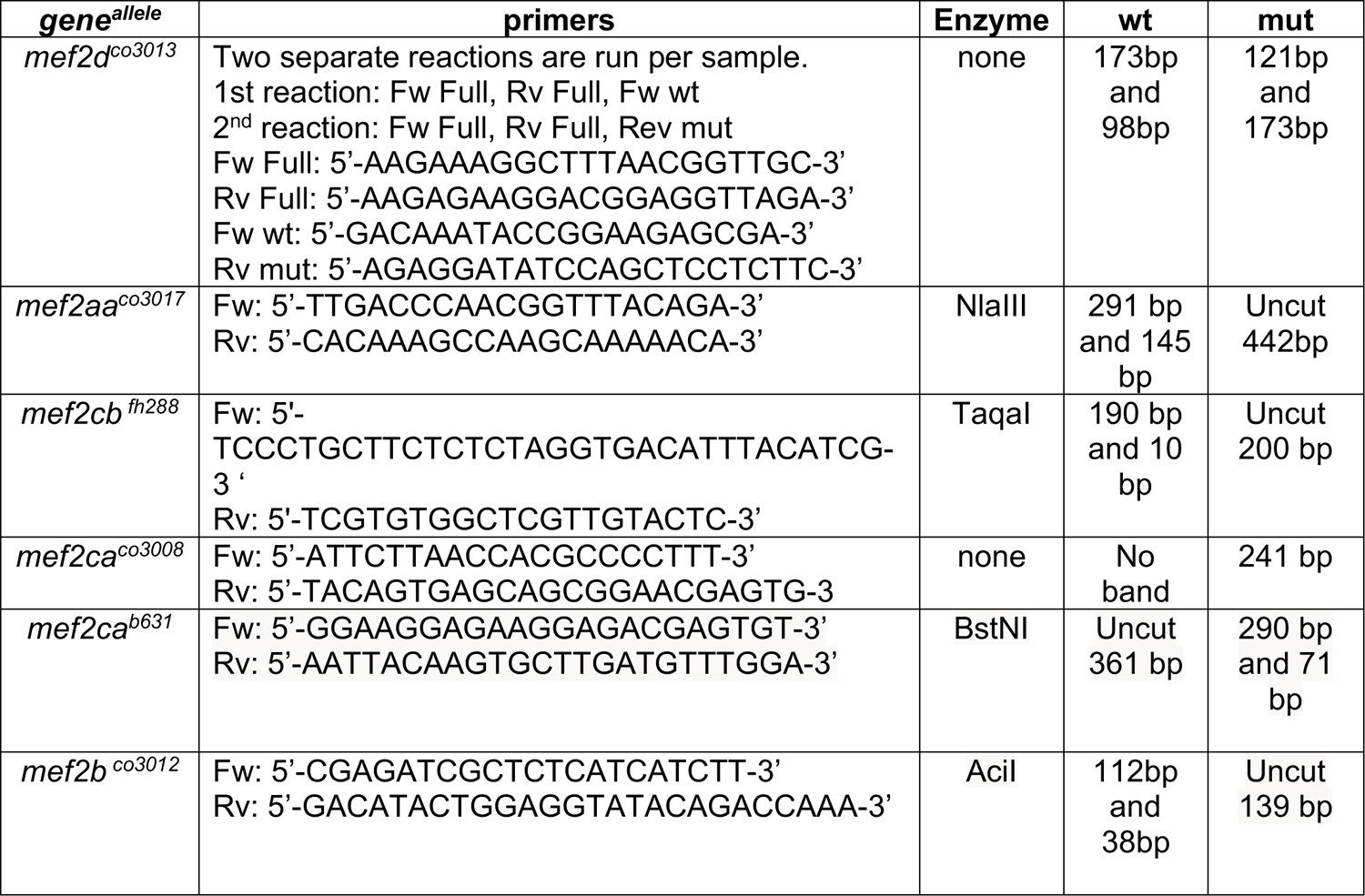

### Sequence Alignments

We used the Clustal Omega tool by EMBL-EBI for multiple sequence alignment of the different paralog gene products. We obtained the following transcripts and protein products from ENSEMBL for alignments: mef2aa-206 ENSDART00000171594.2, mef2ab-201 ENSDART00000173414.2, mef2b-202 ENSDART00000166300.3, mef2ca-202 ENSDART00000099134.5, mef2cb-207 ENSDART00000183585.1, mef2d-203 ENSDART00000132589.2ENSDART00000132589.2

### Statistical analyses

Penetrance scores were compared using Fisher’s exact test to determine significance. All scoring data and exact p values are reported in supplementary data table. Studies with symplectic cartilage measurements are presented as box and whisker plots, the box extends from the 25th to 75th percentiles. The line in the middle of the box is plotted at the median and the bars are minimum and maximum values. We used a Dunnet’s T3 test to compare total symplectic cartilage (left plus right sides) between genotypes. For developmental instability, the absolute value of the difference between left and right symplectic cartilage lengths were grouped by genotype, and Brown-Forsythe and Welch ANOVA tests were used to determine significant differences in left-right asymmetry between genotypes. Power analyses were used to determine the number of animals to be examined for each experiment. All statistical analyses and exact p-values are reported in supplementary data table.

## Ethics statement

All of our work with zebrafish has been approved by the University of Colorado Institutional Animal Care and Use Committee (IACUC), Protocol # 00188. Animals were euthanized by hypothermic shock followed by 1.5% sodium hypochlorite.

## Materials availability statement

All of the materials created for this study are freely available upon request without restrictions.

## Acknowledgements

We thank Charles Kimmel for careful reading of this manuscript, members of the department of craniofacial biology for insightful discussions, Austin Tillery for contributions in the early stages of this study, and the University of Colorado zebrafish care staff. This work was supported by the National Institutes of Health [R01 DE029193 to J.T.N. and F32 DE029995 to J.M.M.] and the National Science Foundation [GRFP 201569 to R.B.Z].

